# High neutralizing antibody levels against SARS-CoV-2 Omicron BA.1 and BA.2 after UB-612 vaccine booster

**DOI:** 10.1101/2022.03.18.484436

**Authors:** Farshad Guirakhoo, Shixia Wang, Chang Yi Wang, Hui-Kai Kuo, Wen-Jiun Peng, Hope Liu, Lixia Wang, Marina Johnson, Adam Hunt, Mei Mei Hu, Thomas P. Monath, Alexander Rumyantsev, David Goldblatt

**Affiliations:** Vaxxinity Inc., Dallas, TX, USA; United Biomedical Inc Asia; Taipei, Taiwan; Great Ormond Street Institute of Child Health, University College London; London, UK

**Keywords:** SARS-CoV-2, COVID-19, Omicron, variant, vaccine, booster, spike protein, receptor binding domain, antibody, neutralizing antibody

## Abstract

The highly transmissible Omicron variant has caused high rates of breakthrough infections among vaccinated and convalescent individuals. Here, we demonstrate that a booster dose of UB-612 vaccine candidate delivered 7-9 months after primary vaccination increases neutralizing antibody levels by 131-, 61- and 49-fold against ancestral SARS-CoV-2, Omicron BA.1 and BA.2 variants, respectively. Based on the RBD protein binding antibody responses, we estimated a ∼95% efficacy against symptomatic COVID-19 caused by the ancestral strain after a UB-612 booster. Our results support UB-612 vaccine as a potent booster against current and emerging SARS-CoV-2 variants.

## Introduction

In November 2021, the Omicron (B.1.1.529) Variant of Concern (VOC) was first reported in South Africa and quickly spread, becoming the dominant SARS-CoV-2 variant worldwide. Omicron’s high transmissibility and potential for immune system evasion, as suggested by its ability to infect and be transmitted by previously infected and vaccinated individuals, predicted a transmission advantage over the Delta variant and the displacement of the latter as dominant variant [1]. The Omicron variant has 3 major sublineages: BA.1, BA.2, and BA.3. While BA.1 caused most of the cases globally throughout November 2021 and February 2022, BA.2 is now the predominant SARS-CoV-2 variant[2].

The Omicron variant has over 50 new amino acid substitutions with >15 in the receptor-binding domain (RBD) of Spike (S) protein [3]. Although BA.2 shares many mutations with BA.1, these 2 sublineages differ by dozens of amino acid substitutions, especially at the S protein (**Fig. 1A**), some of which could be responsible for the rapid surge in BA.2 cases.

**Figure 1.**
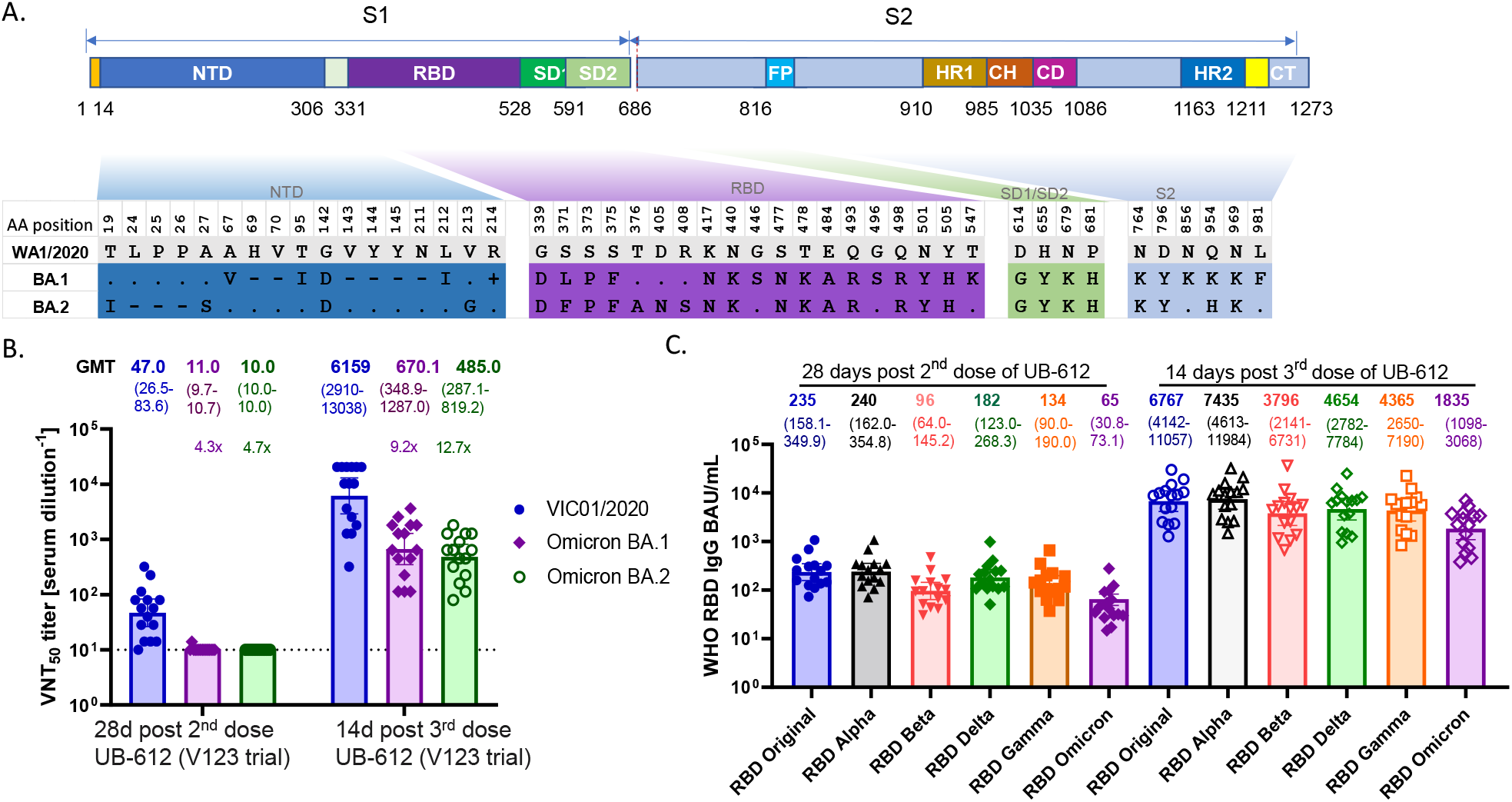
SARS-CoV-2 Omicron BA.1 and BA.2 amino acid substitutions and UB-612 vaccine-induced antibody responses. Panel **A** shows the amino acid substitutions in Omicron’s BA.1 and BA.2 sublineage spike protein. The upper part is the S protein diagram, and the lower part shows the substitutions. The “.”, “-” and “+” represent sequence identical, deletion, and insertion in Omicron BA.1 and BA.2 compared with the US-WA1/2020 virus, respectively. Panel **B** shows the GMT VNT_50_ neutralizing antibody titers against SARS-CoV-2 ancestral strain Victoria/1/2020 (VIC01/2020) and Omicron (B.1.1.529) variant sublineages BA.1 and BA.2 in sera from Phase 1 trial (V123) participants (n=15). The sera were collected at 28 days after 2 doses and at 14 days after the booster dose with UB-612 (100 µg). Data expressed in the reciprocal dilutions for each serum sample and GMT (95% CI) are plotted. Panel C shows the RBD-specific IgG binding titers against SARS-CoV-2 major VOCs in sera collected 28 days after 2 doses and 14 days after 3 doses with UB-612 (100 µg) from Phase 1 trial participants (n=15). The loss of antibody binding to the RBD of variants compared with the original RBD (ancestral strain) remains stable between 2 and 3 doses of UB-612 vaccine, despite an increase in levels of binding antibodies to RBD. GMT, geometric mean titers; VNT, virus neutralization test.

Given that over 90% of neutralizing antibodies are present in plasma of convalescent individuals and up to 99% of neutralizing antibodies elicited by vaccination with the mRNA-1273 vaccine are directed to RBD [4], these mutations could be largely responsible for Omicron’s ability to evade neutralizing antibodies induced by the approved COVID-19 vaccines [5]. Multiple studies have shown a 20-to 30-fold reduction in neutralizing antibody activity against Omicron in the sera of primary vaccine recipients, compared with the ancestral SARS-CoV-2 or D614G viruses [5–7]. The emergence of new variants, in addition to rapidly waning immunity over time, has raised concerns about breakthrough infections in vaccinated individuals, and highlights the need for booster doses. Homologous or heterologous booster vaccines, all based on full-length S protein, restored protective neutralizing antibodies to levels achieved by the primary immunization; however, these titers were 7.1-fold lower against Omicron BA.1 than the ancestral strain, suggesting a continued risk of breakthrough infections in vaccinated individuals over time [5].

In contrast to most of approved COVID-19 vaccines, which encode the full-length S protein, the UB-612 vaccine candidate is composed of Wuhan-Hu S1-RBD-sFc fusion protein and is enriched with 5 peptides representing Sarbecovirus-conserved Th and CTL epitopes on the S2 subunit, Membrane (M), and Nucleocapsid (N) proteins [8]. A favorable safety and tolerability profile for UB-612 was demonstrated in ∼4000 participants in a Phase 1 trial and a Phase 2 trial conducted in Taiwan [9]. In both trials, the UB-612 vaccine was found to have a favorable safety profile and low reactogenicity. Two immunizations with UB-612 led to a >90% seroconversion rate of neutralizing antibodies in vaccine recipients. In these same studies, UB-612 was shown to elicit long-lasting neutralizing antibody titers similar to levels detected in convalescent patients [10], that were cross-reactive against Delta and Omicron variants [9].

The objectives of this study were to evaluate the magnitude of neutralizing antibodies elicited by a third dose (booster) of UB-612 against Omicron and their reactivity to recombinant S and RBD protein antigens across various variants.

## Methods

After receiving a 2-dose primary vaccine series or a booster given 7-9 months after the second dose, sera from 15 participants (who consented to be in this study) in the Phase 1 trial (UB-612, 100-µg dose), were tested in a SARS-CoV-2 live virus neutralization test (VNT) at Vismederi, Siena, Italy (a Coalition for Epidemic Preparedness Innovations (CEPI) testing laboratory for COVID-19 vaccines) **(Table S1)**. Previously, to establish an International Reference Standard for anti−SARS-CoV-2 antibody detection, the VNT used in our analysis and performed by Vismederi, was compared with other VNTs and found to be the most stringent assay, resulting in a lower geometric mean titer (GMT) than other plaque reduction-, foci reduction-, cytopathic effect (CPE)-, or pseudotyped virus-based neutralization assays [11]. The detailed materials and methods for clinical trial and sample information, neutralization assays, immunological assays and statistical analyses are described in the Supplemental Materials.

## Results

### A booster dose of UB-612 significantly increased the levels of neutralizing antibodies against the ancestral strain and Omicron variants

Two doses of UB-612 showed modest neutralizing activity against the authentic wild-type SARS-CoV-2 ancestral strain (Victoria/1/2020) (GMT VNT_50_ of 47.0), and very low activity against Omicron’s BA.1 and BA.2 sublineages (GMT VNT_50_ of 10-11) (**Fig. 1B**) (n=15). Similarly, a 2-dose immunization with mRNA vaccines has been shown to result in low levels of Omicron neutralizing antibody responses: (i) mRNA-1273 on day 21 after immunization stimulated a GMT pVNT_50_ of 14, and (ii) BNT162b2 on day 28 after immunization lead to a GMT pVNT_50_ of 7 [6,7].

A booster dose of UB-612 delivered at 7-9 months after the primary series increased neutralizing antibody titers against the ancestral strain, as well as Omicron BA.1 and BA.2 variants, to GMT VNT_50_ of 6,159, 670, and 485, respectively, which constitutes 131-, 61-, and 49-fold higher GMTs than those achieved after 2 doses **(Fig. 1B)**. The estimated decrease in neutralization titers against Omicron BA.1 and BA.2 in 15 sera obtained 2 weeks after UB-612 booster, was 9.2- and 12.7-fold, respectively, compared with the ancestral Victoria strain. Previously, only a 5.5-fold decrease against BA.1 was reported when 20 sera from UB-612 vaccinated participants were tested in a pseudovirus-based neutralization assay (GMT pVNT_50_ of 12,778 against the Wuhan strain, vs. 2,325 against the BA.1 strain) [9]. These data support the breadth of UB-612−elicited neutralizing antibodies across multiple SARS-CoV-2 variants after the booster, a differentiation property of UB-612 primarily attributed to its subunit protein RBD antigenic component [10].

### Expansion of antibody breadth following UB-612 booster

We evaluated reactivity of UB-612−elicited antibodies to S and RBD proteins, using 15 sera from the Phase 1 trial (V123) participants and 84 randomly selected sera from Phase 2 trial participants immunized with UB-612 (**Tables S1-S2**). These sera were tested in 2 ELISA-based assays for IgG direct binding to recombinant S and RBD proteins, and inhibition of recombinant S and RBD protein binding to the human angiotensin-converting enzyme 2 (hACE2) receptor. A third dose of UB-612 booster immunization stimulated broadly reactive IgG antibodies, effectively binding to RBDs of 14 divergent SARS-CoV-2 variants, including Alpha, Beta, Gamma, Delta, and Omicron (**Fig. 1C** and **Fig. S1**).

Compared with the second UB-612 dose, IgG binding titers against Omicron’s RBD after the third dose increased by over 40-fold, and the titers against RBDs of other SARS-CoV-2 variants were also increased 30- to 50-fold after the booster dose. When the IgG titer ratio (in binding antibody units (BAU)/mL) of several variants was compared to the ancestral Wuhan strain, the normalized RBD antibody-binding responses to the tested variants were found to be similar after 2 or 3 doses: Alpha (0.98-fold), Beta (2.44-fold), Delta (1.33-fold), Gamma (1.77-fold), and Omicron (3.3-fold) after 2 doses; and Alpha (0.91-fold), Beta (1.8-fold), Delta (1.4-fold), Gamma (1.55-fold), and Omicron (3.7-fold) after 3 doses.

Similar to RBD binding, the results of the S-protein binding antibody responses (S:ACE2- and RBD:ACE2-blocking antibody titers), confirmed the extent of stability in ratios of parental to variant IgG antibodies stimulated by 2 or 3 doses of UB-612, despite an up to 60-fold increase in titers against different variants after the booster dose (**Fig. S2, Fig. S3**).

### Vaccine efficacy prediction

We compared the level of UB-612−elicited IgG antibodies with data previously reported for several authorized vaccines determined in equivalent S- and RBD-binding assays [12]. After a 2-dose primary immunization series, the GMTs of UB-612−elicited IgG antibodies were 69 and 127 (BAU/mL) against the Wuhan S protein, and 235 and 494 (BAU/mL) against the RBD antigen in sera from Phase 1 and Phase 2 participants, respectively (**Fig. S4**). These IgG responses were comparable to those observed in individuals after the primary immunization with adenovirus vectored vaccines (1-dose Ad26.COV2.S or 2-dose ChAdOx1-S) but were lower than the response observed after 2 immunizations with mRNA vaccines. The additional booster dose with UB-612 increased levels of both S- and RBD-protein binding IgG antibodies in the Phase 1 participants by more than 16- and 13-fold, and increased antibody GMTs to 2,138 and 6,767 (BAU/mL), respectively, matching those achieved by 2 immunizations with the mRNA vaccines.

We further utilized a vaccine efficacy prediction model based on the RBD activity of IgG antibodies to the ancestral strain, extending previous models based on neutralizing antibodies [13] or S protein−binding activities [12]. According to this model, the predicted vaccine efficacy of UB-612 against symptomatic disease caused by the prototype strain after 2 doses is ∼72% (235 BAU/mL) with sera from 15 Phase 1 participants, ∼82% (494 BAU/mL) with sera from 84 randomly selected Phase 2 participants, and ∼95% after the booster dose (6,767 BAU/mL), with sera from 15 Phase 1 participants (**Fig. 2 and Fig. S4**).

**Figure 2.**
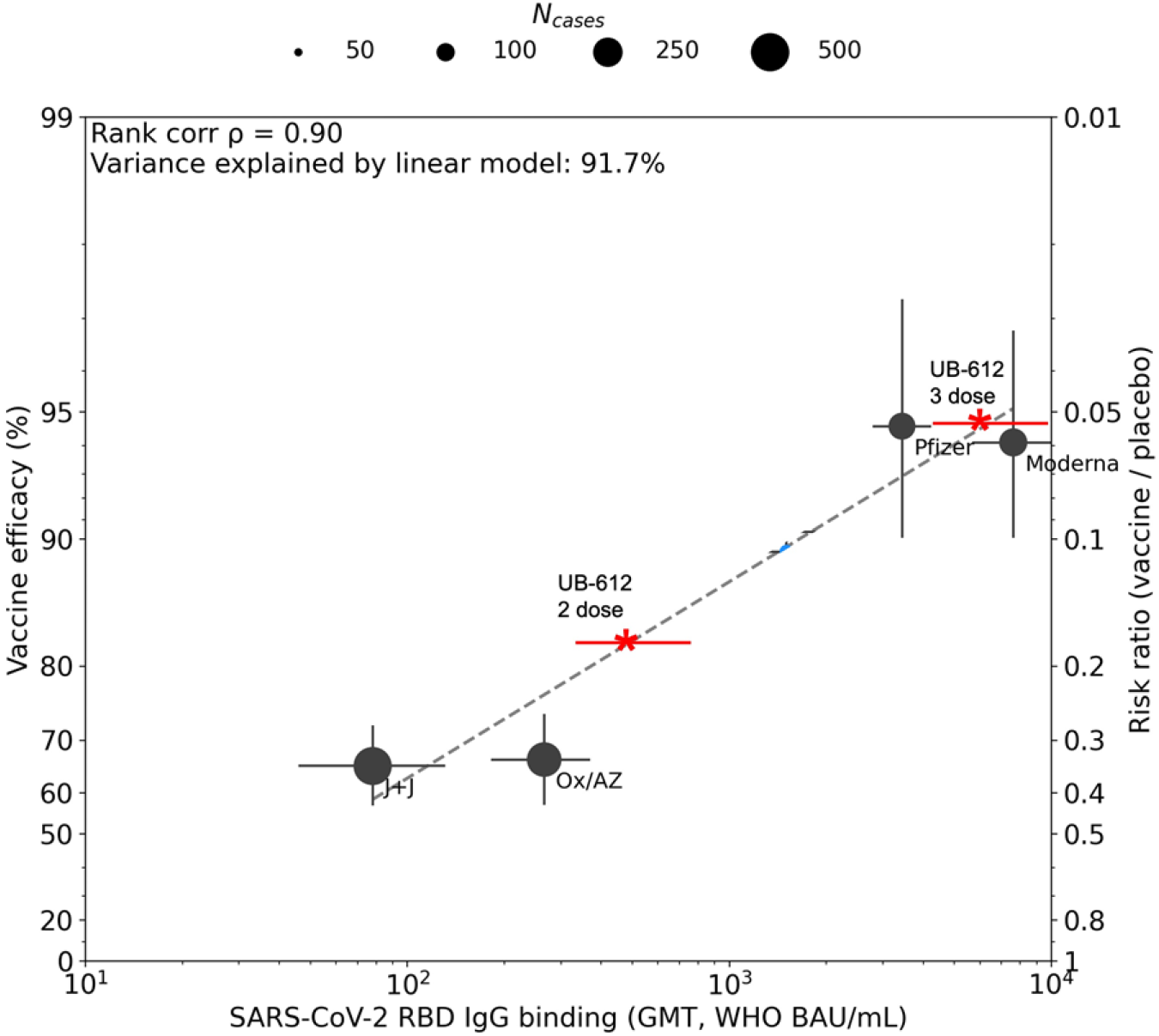
Estimated UB-612 efficacy after 2 and 3 doses of UB-612 vaccine. A model bridging vaccine-induced RBD IgG response to vaccine efficacy against symptomatic COVID-19 caused by ancestral Wuhan [5]. Estimated efficacy of UB-612 after 2 doses is ∼72% (CI, 70%-80%) based on RBD binding IgG antibodies from 15 participants (Phase 1) (GMT 235 BAU/mL, 95% CI, 158-350, ∼82% (CI, 80%-85%) based on RBD binding IgG antibodies (GMT 494 BAU/mL, 95% CI, 337-725, shown in this graph), and ∼95% (93%-97%) after a booster vaccination (GMT 6767 (95% CI, 4142-11,057). GMT, geometric mean titers.

## Discussions

It is difficult to compare neutralization activities of different vaccine platforms against variants because there are currently no international standard reagents available for variants, and each manufacturer uses a different assay, including live virus, pseudovirus-based, or chimeric recombinant virus, with different endpoint readouts, such as colorimetric or CPE-based evaluations. Moreover, each stock virus may contain different ratios of noninfectious to infectious particles and different virus concentrations may be used for neutralizing sera from vaccinees, both of which could influence the resulting titers. Keeping these caveats in mind, we observed GMT levels after the UB-612 booster dose comparable to those elicited by a booster dose of mRNA vaccine BNT162b2 [7] and mRNA-1273-50 µg [6]. After the booster dose (third vaccination), the live virus neutralizing antibody GMTs against the ancestral strain and Omicron BA.1 were 763 and 106 for BNT162b2 (7.2-fold loss) [7], and 2,423 and 850 for mRNA-1273- 50 µg (2.8-fold loss) [6]. The pseudovirus neutralization GMT titers were 6,539, 1,066, and 776 against the ancestral strain WA1/2020, Omicron BA.1, and BA.2, respectively, at 14 days after the third dose of BNT162b2 (a 6.1- and 8.4-fold loss against BA.1 and BA.2, respectively, compared with the ancestral strain). The homologous booster of mRNA-1273, BNT162b2, or Ad26.COV.S S-based vaccines dramatically increased neutralizing antibodies to Omicron (20- to 30-fold) compared with the modest increase reported for the ancestral strain (1- to 4-fold), likely due to a higher baseline titer for the ancestral strain compared with the variants.

Vaccination with UB-612 elicited highly cross-reactive IgG and neutralizing antibodies to Omicron variants (with a 49- to 61-fold increase in VNT_50_) and the ratio of binding antibodies to ancestral strain/Omicron and other variants remained stable after the second and booster immunizations. It has been demonstrated that the third dose of mRNA vaccines containing the full-length S protein of SARS-CoV-2 can recall the persisting memory B cells clones that have been affinity-matured after a long interval between the doses, as well as to develop new clones of memory B cells to produce neutralizing antibodies to conserved RBD regions, enhancing the breadth of cross-variant neutralization [14]. We believe that the UB-612 vaccine, which contains only the RBD region of the S protein, may be more effective in recalling persisting memory B-cell clones against the conserved epitopes to protect against variant-induced serious disease. Another potential advantage of the UB-612 vaccine is the inclusion of Th/CTL peptides from S2, M and N proteins that are highly conserved in Omicron variants (data not shown). There is growing evidence supportive of importance of vaccine-elicited CD8+ T cells in protection against Omicron [15].

In summary, a booster dose of UB-612 elicited robust S- and RBD-specific binding and virus neutralizing antibodies against multiple SARS-CoV-2 variants, including Omicron BA.1 and BA.2. The magnitude and extent of reactivity of the neutralizing antibody responses after UB-612 booster match those reported for the authorized vaccines, including BNT162b2 and mRNA-1273. Additionally, UB-612 has been shown to stimulate T-cell responses against conserved S2, N, and M peptides [9,10] and may provide long-lasting antibody responses [11] that would further differentiate UB-612 from many authorized vaccines. As SARS-CoV-2 continues to evolve, several strategies are being explored to effectively prevent COVID-19 caused by newly emerging SARS-CoV-2 variants, including monovalent variant antigen matching, multivalent, or universal vaccine approaches. Our results indicate that UB-612 could offer an alternative strategy for a booster vaccine, eliciting long lasting neutralizing antibody responses with extensive activity across currently circulating and potentially future SARS-CoV-2 variants.

## Acknowledgments

We thank many colleagues at UBIAsia, UBP, and Vaxxinity who designed, developed, and produced the UB-612 vaccine candidate and performed the Phase 1 and 2 clinical trials in Taiwan. We also thank the Phase 1 (NCT04545749) and Phase 2 (NCT04773067) clinical trial participants, from whom the post-vaccination sera were obtained. We are also grateful to the Vismederi team for their work on live virus-neutralizing antibody assays.

## Author contributions

Conceived and conceptualized, wrote the manuscript: FG

Analyzed data, interpreted data, wrote the manuscript: SW

Received consent from the patients for the samples to be used for this study and organized shipment of their sera for binding and neutralization assays and reviewed the manuscript: CYW, WJP, HL, HKK

Assisted in planning of the study: MMH, TPM

Wrote and reviewed the manuscript: AR

Analyzed data: LW

Performed experiments and interpreted data: DG, MG, AH

## Previous presentation of data

Part of data was previously presented at the World Vaccine Congress, April 17-20, 2022; and was published in BioRxiv.

