## Supplemental Materials for "High neutralizing antibody levels against SARS-CoV-2 Omicron BA.1 and BA.2 after UB-612 vaccine booster"

Figure S3. ACE2 binding blocking antibody titers in vaccinated participants at 30 days post 2 doses (2x UB-612) and at 14 days post 3 doses (3x UB-612) of UB-612 vaccination (n=15). 13
Figure S4. RBD binding IgG (Panel A) and spike binding IgG (Panel B) against SARS-CoV-2 original Wuhan isolate in vaccinated participants at 30 days post 2 doses (2x UB-612) and at 14 days post 3 doses (3x UB-612) of UB-612 vaccination from the Phase 1 study (V123 study) (n=15) (see Table S1 and S2). ........ 14

**Materials and Methods**

Clinical trials

Phase 1 (NCT04545749) was an open-label trial to evaluate safety, tolerability, and immunogenicity of ascending doses (10, 30, and 100 μg) of UB-612 given by injection to healthy adults aged 20-55 years (n=60, 20 participants in each dose arm) in 2 doses with a 28-day interval. Follow-up of participants for safety and immunogenicity was conducted for 6 months, with blood draws performed on days 14, 28, 42, 56, 112, and 196 after the first dose. A separate extension study (NCT04967742) was conducted in participants (n=50, including 18 in the 100-μg UB-612 dose arm) who completed 2 vaccinations, where an additional 100 μg UB-612 third dose booster was delivered 6 months after the second dose of the primary series. The participants were followed up for safety and immunogenicity, with blood samples collected on days 1, 14, and 84 post booster.

Phase 2 (NCT04773067) is an ongoing observer-blind, multiple-center, randomized, placebo-controlled study evaluating the immunogenicity, safety, tolerability, and lot consistency of 2 doses of UB-612 vaccine in participants aged 12-85 years in Taiwan. All adult participants were randomly allocated to receive 2 doses of 100 μg UB-612 or normal saline as placebo in a 6:1 ratio (n=3850). At least 350 adults (aged >18 to <65 years old) and 154 elderly (aged ≥65 years) were assessed for immunogenicity, with blood collected on days 1, 29, 57, and 197.

Serum specimens

Of the18 participants in the Phase 1 study, 15 elected to participate in the Phase 1 extension booster study (for a total of 3 100 μg UB-612 doses received across Phase 1 studies) and consented to additional testing of their blood samples collected on days 28 and 14 after the second and third immunizations, respectively.

From the Phase 2 immunogenicity cohort, a total of 92 samples collected on day 28 after the second injection were randomly selected for additional antibody testing. Eighty samples were selected from the UB-612 study arm and 9 from the placebo control arm.

### Cell culture

Vero E6 cells (American Type Culture Collection [ATCC, #CRL-1586]) were cultured in Dulbecco's Modified Eagle's Medium (DMEM) high glucose supplemented with 2 mM L-glutamine, 100 U/mL of penicillin-streptomycin mixture (complete DMEM) and 10% fetal bovine serum (FBS). Cells were maintained at 37°C, 5% CO2 and passaged every 3-4 days. The day prior to the execution of the microneutralization assay, VERO E6 cells diluted in complete DMEM 2% FBS were added to 96-well plates (1.5 x $10^4$ cell/well) and incubated at 37°C, 5% $CO_2$ until use.

### Viruses

Authentic Wild Type SARS CoV-2 2019 (Victoria/1/2020 strain) was kindly provided by Coalition for Epidemic Preparedness Innovations (CEPI) and Omicron SARS-CoV-2 variant 1.1.529, subvariant BA.1, and subvariant BA.2 were isolated by Vismederi using combined oropharyngeal/nasal procedure and the sequences were verified. Spike mutations in the Omicron subvariant BA.1: A67V - H69del - V70del - T95I - G142D - V143del - V144del - Y145del - N211del - L212I - G339D - S371L - S373P - S375F - K417N - N440K - G446S - S477N - T478K - E484A - Q493R - G496S - Q498R - N501Y - Y505H - T547K - D614G - H655Y -

N679K - P681H  R682G - N764K - D796Y - N856K - Q954H- N969K - L981F); spike mutations in Omicron subvariant BA.2:  T19I - L24del - P25del - P26del - A27S - V213G - G339D - S371F - S373P - S375F - T376A - D405N - R408S - K417N - S477N - T478K - E484A - Q493R - Q498R - N501Y - Y505H - D614G - H655Y - N679K - P681H - R682W - N764K - D796Y - Q954H - N969K.

The viruses were propagated as described elsewhere [1]. Briefly, viral propagation was performed in 175 $cm^2$ tissue-culture flasks pre-seeded with 50 mL of VERO E6 cells (1 x $10^6$ cells/mL) diluted in DMEM 10% FBS. After a 24-hour incubation period at 37°C, 5% CO2, flasks were washed twice with sterile Dulbecco's phosphate buffered saline (DPBS) and then inoculated with the SARS-CoV-2 virus. The sub-confluent cell monolayer was allowed to incubate with the virus for 1 hour at 37°C, 5% $CO_2$, and flasks were then filled with 50 mL of DMEM 2% FBS and incubated at 37°C, 5% $CO_2$. Cells were monitored daily until manifestation of an 80%-90% cytopathic effect (CPE). Supernatants of the infected cultures were then harvested, centrifuged at 1000 rpm for 5 minutes at 4°C to remove cell debris, aliquoted, and stored at -80°C.

The propagated viral stocks were titrated in 96-well plates containing a sub-confluent VERO E6 cell monolayer. A 10-fold serial dilution of virus ($10^{-1}$ to $10^{-11}$) was incubated with the cells and checked daily for signs of CPE for a total of 3 days (Wild Type) or 4 days (Omicron BA.1 and BA.2 variants). The viral titer was calculated using the 50% tissue culture infectious dose per mL ($TCID_{50}$/mL) as the endpoint and defined as the reciprocal of the highest virus dilution yielding at least 50% CPE in the inoculated wells according to the Reed and Muench formula [2].

SARS-CoV-2 live-virus neutralization assay

SARS-CoV-2 live-virus neutralization was performed via a microneutralization assay based on CPE at Vismederi S.r.l., Siena, Italy. Briefly, sera were heat inactivated for 30 minutes at 56°C, serially diluted in DMEM 2% FBS (at 2-fold; starting dilution: 1:10), and added in duplicate to 2 different 96-well plates. The plates were then incubated for 1 hour at 37°C, 5% CO2, with 25 median tissue culture infectious dose ($TCID_{50}$) of authentic SARS-CoV-2 Wild Type virus strain Victoria/1/2020 or Omicron variants (serum-virus ratio 1:1) to allow binding of antigen-specific antibodies to the virus. The virus-serum mixture was then added to sub-confluent VERO E6 cells. After 3 days (Wild Type) or 4 days (Omicron), cells were inspected for signs of CPE under an inverted light microscope. The reciprocal of the highest sample dilution able to protect at least 50% of the cells from CPE was regarded as the neutralization titer and reported as geometric mean titer (GMT) (mean of 2 replicates). In the absence of neutralization, an arbitrary titer value of 5 (half of the limit of detection [LOD]) was reported. GMT values equal to 5120 (which corresponds to the assay upper limit of detection [ULOD]) were reported as the next dilution up (10,240) for statistical and graphical purposes. The First WHO International Standard panel for anti−SARS-CoV-2 immunoglobulin, human (code 20/268) developed by the National Institute for Biological Standards and Controls (NIBSC) was used as the Reference Panel in each assay.

Immunological assays

Samples were analyzed for the presence of immunoglobulin G (IgG) in SARS-CoV-2 nucleocapsid protein, receptor binding domain (RBD) of S1, and trimeric spike antigen derived from the original Wild Type Wuhan strain as previously described [3], as well as spike and RBD responses to Alpha, Beta, Gamma, and Delta strains (MSD® SARS-Coronavirus Plate 7, Rockville, MD) at the University College London. The MSD IgG assay was calibrated against the WHO international anti−SARS-Cov-2 antibody standard, which assigns a value of 1000 binding antibody units (BAU) for the spike, RBD, and nucleocapsid antigens. The lower limits of quantitation for Wild Type spike and RBD were 1.09 and 145 BAU/mL, respectively. As no standards exist for the variants, the internal standard used was evaluated and adjusted for the variant antigens based on the binding signal obtained. The MSD assay was also used to evaluate ACE2 receptor inhibition against the relevant antigens highlighted above as previously described [3].

### Statistical analysis

Descriptive statistics were summarized for GMT and 95% CI for the GMT was calculated.

**<u>Table S1.</u> Summary of Demographics and Other Baseline Characteristics of Participants from UB-612 Phase 1 Open-label Extension Study (NCT04967742).**

| Demographic Parameters | | UB-621 two 100 μg-doses + booster dose (100 μg)* (N=18) |
|---|---|---|
| Age (years) | n | 18 |
| | Mean | 35.9 |
| | SD | 10.10 |
| | Median | 38.0 |
| | Minimum | 22 |
| | Maximum | 52 |
| Sex | Female | 13 (72.2%) |
| | Male | 5 (27.8%) |
| Ethnicity | Taiwanese | 18 (100%) |
| | Other | 0 |

*Only 15/18 participants who received the 100-ug booster dose consented for their sera to be tested in this study.

**Table S2. Summary of Demographic Characteristics of Randomly Selected Participants From the UB-612 Phase 2 Trial (NCT04773067).**

| Demographic Parameters | | Active | Placebo |
|---|---|---|---|
| Age (years) | n | 84 | 8 |
| | Mean | 52.4 | 55.3 |
| | SD | 18.29 | 18.2 |
| | Median | 62.0 | 63.0 |
| | Minimum | 20 | 24 |
| | Maximum | 79 | 71 |
| Sex | Female | 42 (50%) | 4 (50%) |
| | Male | 42 (50%) | 4 (50%) |
| Ethnicity | Taiwanese | 84 (100%) | 8 (100%) |
| | Other | 0 | 0 |

### Supplemental Figures

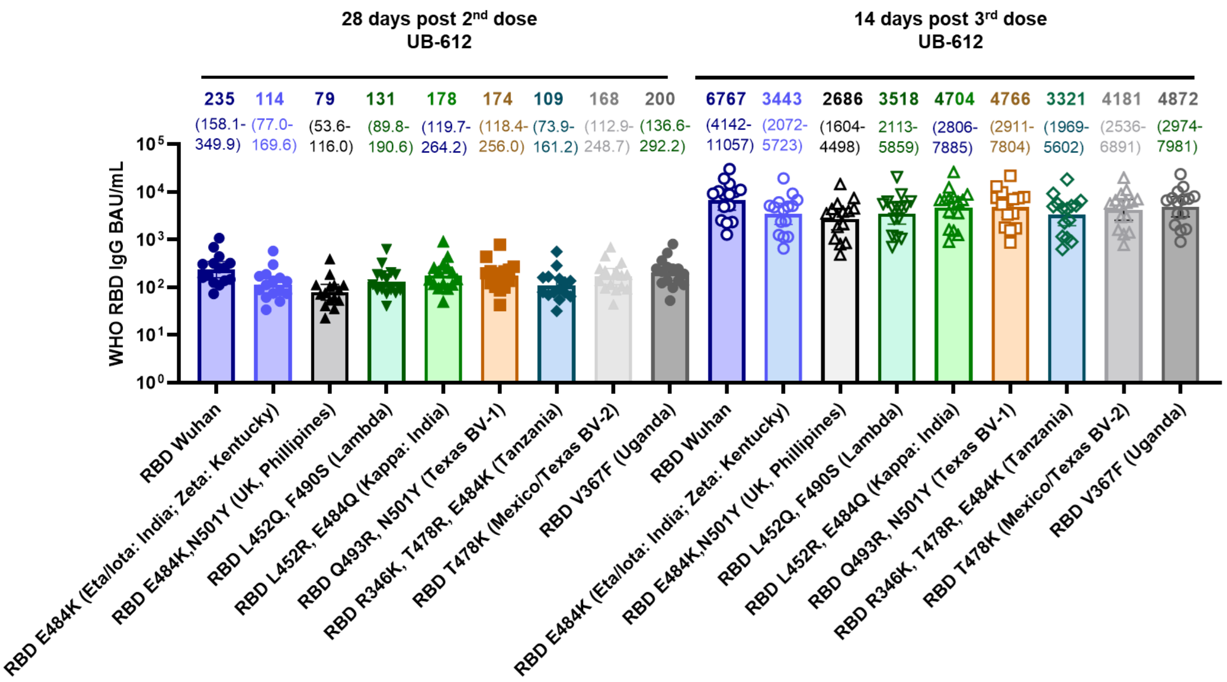

**<u>Fig S1.</u> IgG binding titers against SARS-CoV-2 variants in individuals 28 days post dose 2 (2x UB-612) and 14 days post dose 3 (3x UB-612) of UB-612 vaccination.**

Loss of binding antibodies to RBD for the variants compared with the original RBD (ancestral strain) remains stable between 2 and 3 doses of UB-612 vaccine despite a high increase in levels of binding antibodies to RBD. The ratios of binding antibodies against original RBD vs. variants (from left to right, e.g., RBD E484K to RBD V367F) are 2.0, 3.0, 1.8, 1.3, 1.3, 2.1, 1.4, and 1.2 for 2 doses, respectively; and 2.0, 2.5, 1.9, 1.4, 1.4, 2.0, 1.6, and 1.4 for 3 doses, respectively. Numbers above each bar represent GMT and 95% CI.

GMT, geometric mean titer; RBD, receptor-binding domain.

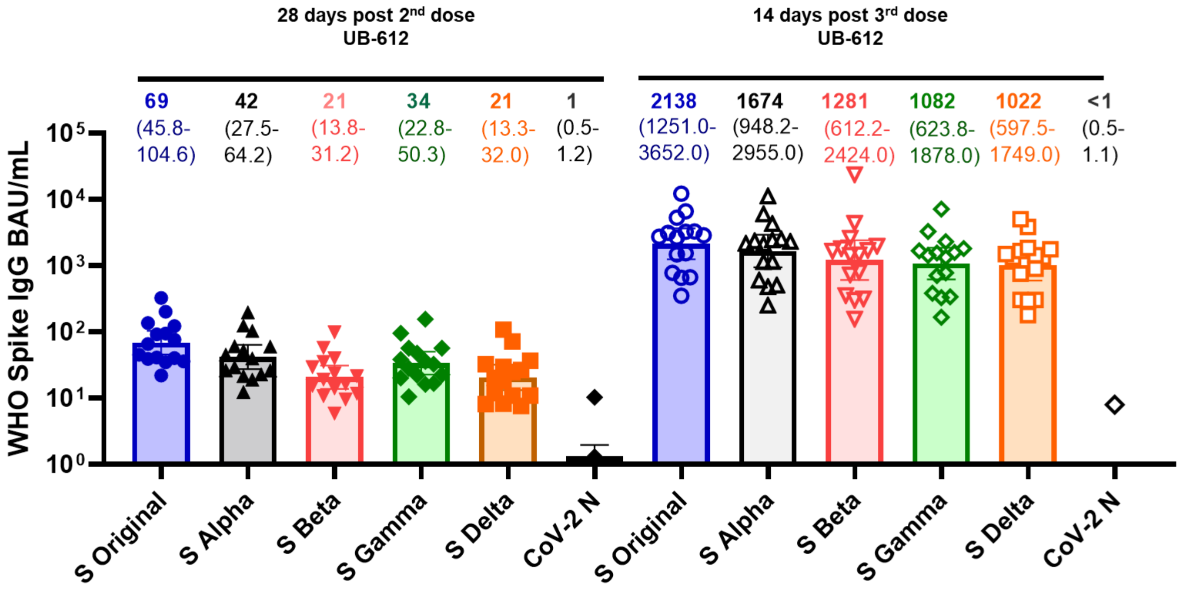

**Fig S2. Spike protein-specific binding IgG against SARS-CoV-2 major VOCs in the sera of Phase 1 participants (n=15) collected 30 days after the second dose and 14 days after the third dose of UB-612 vaccination.**

Binding to SARS-CoV-2 N protein (CoV-2 N) was used as a control in the binding assay to demonstrate that participants were not naturally infected with SARS-CoV-2 (<10 BAU/mL). After 2 doses, decrease in binding titers to the spike protein of VOC compared with the original (ancestral) spike was 1.6-, 3.3-, 2.0-, and 3.3-fold for Alpha, Beta, Gamma, and Delta, respectively. After a booster dose these numbers remained relatively stable at 1.3-, 1.7-, 2.0-, and 2.1-fold decreases, for Alpha, Beta, Gamma, and Delta, respectively. Numbers above each bar represent GMT and 95% CI.

GMT, geometric mean titer; IgG, immunoglobulin G; VOC, Variant of Concern.

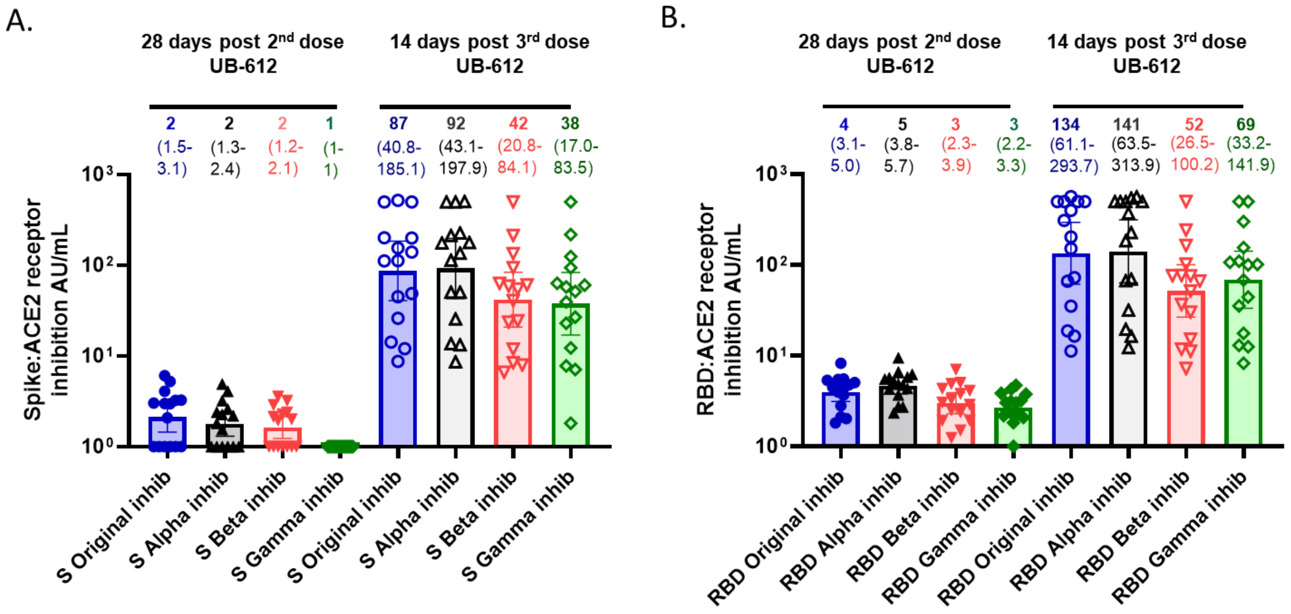

**<u>Fig S3.</u> ACE2 blocking antibody titers in vaccinated participants at 30 days post 2 doses (2x UB-612) and at 14 days post 3 doses (3x UB-612) of UB-612 vaccination (n=15).**

A: spike protein (S):ACE2 blocking Ab against VOCs; B: RBD:ACE2 blocking Ab against VOCs (see Table S1). After 2 doses, the loss in binding inhibition for the spike protein of VOC compared with the original (ancestral) spike is 0-, 2.0-, and 2.0-fold for Alpha, Beta, and Gamma, respectively. After a booster dose these numbers remain relatively stable at 0-, 1.7-, 2.1-, and 2.3-fold for Alpha, Beta, and Gamma, respectively (Panel A). Binding inhibition for the RBD protein of VOC compared with the original (ancestral) spike was 0-, 1.3-, and 1.3-fold for Alpha, Beta, and Gamma, respectively. After a booster dose, these numbers remained relatively stable at 0-, 2.6-, and 1.9-fold for Alpha, Beta, and Gamma, respectively (Panel B). Numbers above each bar represent GMT and 95% CI.

GMT, geometric mean titer; RBD, receptor-binding domain; VOC, Variant of Concern.

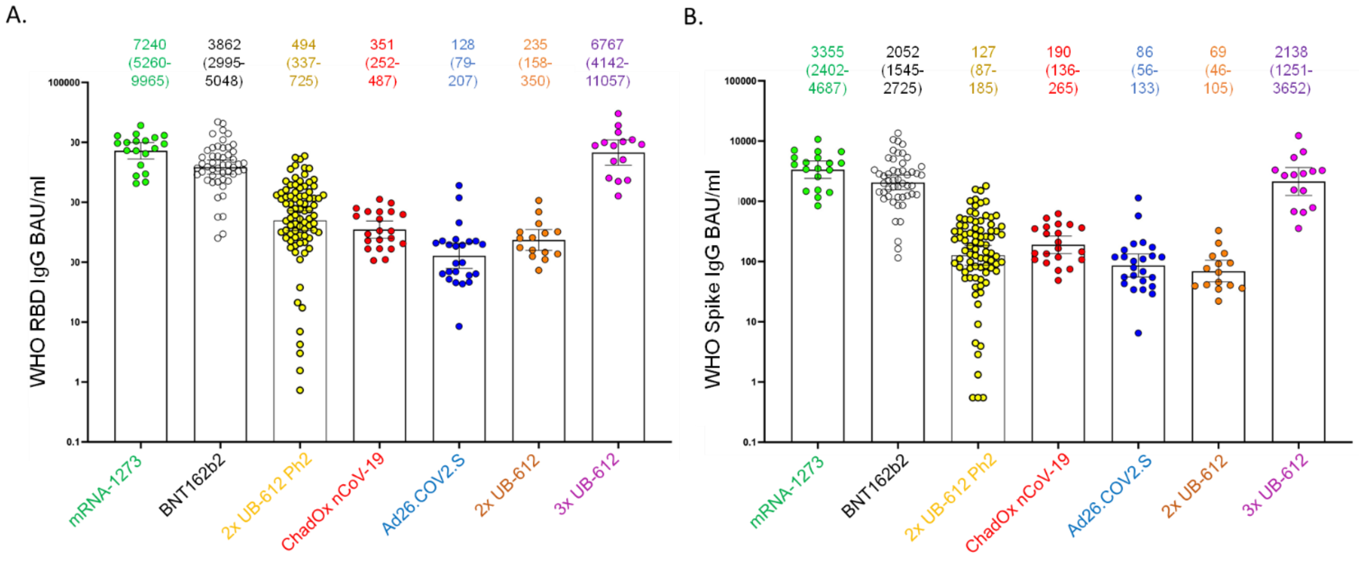

**<u>Fig S4.</u> RBD binding IgG (Panel A) and spike binding IgG (Panel B) against SARS-CoV-2 original Wuhan isolate in vaccinated participants at 30 days post 2 doses (2x UB-612) and at 14 days post 3 doses (3x UB-612) of UB-612 vaccination from the Phase 1 study (V123 study) (n=15) (see Table S1 and S2).**

Sera from a subset of UB-612 vaccinated participants from the Phase 2 trial (V205) (n=84) drawn at 14 days post 2 doses (see Table S1) were also included. For EU-approved vaccines, sera were collected from vaccinated participants (given at 1 or 2 doses) after 7, 8, 8, and 34 days (median) for mRNA1273, BNT162b2, ChadOX1, and Ad26. COCV2.S, respectively. The median time between doses was 3-4 weeks (for ChadOX1 vaccine median time was 66 days and Ad26. COCV2.S was given at a single dose) [4]. Numbers above each bar represent GMT and 95% CI. Note: The GMT numbers in this figure are different than those published in Fig. 1 by Goldblatt et al [5]. This is because the samples described in the Goldblatt et al paper were above the upper limit of detection for the assay, and therefore were assigned arbitrary values. In this figure, those samples were further diluted and retested to determine the exact titers with accurate concentrations at the upper limit of detection. GMT, geometric mean titer.
